# NeoHeadHunter: an algorithm for the detection, ranking and probabilistic classification of neoepitope candidates

**DOI:** 10.1101/2023.10.25.563895

**Authors:** Xiaofei Zhao

## Abstract

**Background:** The manufacturing of personalized cancer vaccine requires the accurate identification of neoepitopes, abnormal peptides presented by cancer cells and recognized by the host immune system of the cancer patient.

**Results:** We designed and developed NeoHeadHunter, a computational algorithm and pipeline to detect and rank neoepitope candidates. Unlike other algorithms, NeoHeadHunter can estimate the probability that each predicted neoepitope candidate is true positive. To evaluate NeoHeadHunter, we used the Tumor neoantigEn SeLection Alliance (TESLA) data-set derived from the sequencing of nine patients and characterized by 44 experimentally validated positive neoepitopes, a data-set derived from the sequencing of three cancer patients and characterized by eight experimentally validated positive neoepitopes and a manually curated data-set consisting of 64 experimentally validated positive neoepitopes. Our evaluation shows that NeoHeadHunter performs the best compared with other algorithms for both detecting and ranking neoepitope candidates and that NeoHeadHunter can accurately predict such probabilities.

**Conclusions:** NeoHeadHunter can increase the effectiveness of personalized cancer vaccine by sensitively detect, accurately rank and probabilistically classify neoepitope candidates. NeoHeadHunter is released under the APACHE-II license at https://github.com/XuegongLab/neoheadhunter for academic use.

## 1 Introduction

Cancer (i.e., malignant tumor) cells are characterized by a lot of diverse genetic mutations. These mutations can result in the generation of abnormal proteins and peptides. Some of these abnormal peptides can be presented by major histocompability complexes (MHCs) to cancer-cell surface and then recognized by the host immune system, and the alerted immune system can then kills these recognized cancer cells. These abnormal proteins and peptides are referred to as neoantigens and neoepitopes, respectively. Neoepitopes are used as the main ingredient of personalized cancer vaccine which is both highly effective against cancer and characterized by tolerable side effect (fig. 1). Unfortunately, the sensitive detection and accurate ranking of neoepitope candidates are computationally challenging. Moreover, physicians and patients are often interested in predicting treatment response, and such prediction requires the estimation of the probability that an abnormal peptide (i.e., neoepitope candidate) is an immunogenic (i.e., positive as validated by multimer-staining assay) neoepitope that can induce the immune system to kill cancer cells. Currently, an algorithm that can estimate such probabilities does not exist yet.

**Figure 1:**
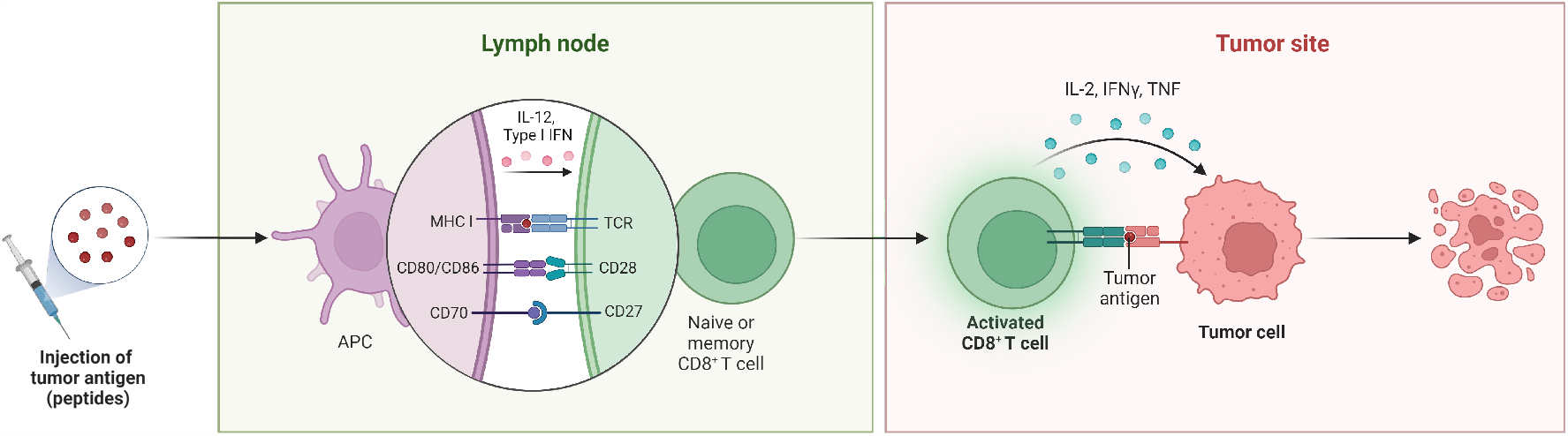
Principle of cancer vaccination by T-cell activation. Adapted from “Cancer Vaccine Principle”, by BioRender.com (2020). Retrieved from https://app.biorender.com/biorender-templates.

We proposed an algorithm named NeoHeadHunter to detect neoepitope candidates and to estimate the probability that each candidate is immunogenic. The key idea of NeoHeadHunter is to use peptideMHC (pMHC) binding affinity, pMHC binding stability, tumor abundance, agretopicity and foreignness (as computed by [Wells et al., 2020]) as features to estimate the probability that each candidate is immunogenic. Wells et al. found a threshold for each of these features so that neoepitope candidates passing all the thresholds for all these features are highly likely to be immunogenic, but it is not clear how to use these features to compute the probability of a candidate being immunogenic. Additionally, NeoHeadHunter is able to detect all immunogenic neoepitopes and to estimate the strength of the immune system of a patient from his/her tumor DNA-seq, normal DNA-seq and tumor RNA-seq data.

## 2 Methods

In essence, NeoHeadHunter uses eq. (12), a logistic regression built upon a decision-tree-like boolean circuit, to rank and probabilistically classify neoepitope candidates. We used the knowledge about the biological mechanism of neoepitopes to propose the general form of eq. (12). We used the six patients with the IDs of 1, 2, 3, 10, 12 and 16 from the TESLA study [Wells et al., 2020] as the discovery cohort to train eq. (12). We used the three patients with the IDs of 4, 8 and 9 from the TESLA study [Wells et al., 2020] as the validation cohort to validate eq. (12). We used the three patients with the IDs of L011, L012 and L013, which were used to evaluate the MuPeXI neoepitope-prioritization algorithm [Bjerregaard et al., 2017], as the independent-validation cohort to test eq. (12) on an independent study. We used the pMHCs manually curated by Koşaloğlu-Yalçın et al. [2018], which were used to evaluate the DeepHLApan immunogenicity-prediction algorithm [Wu et al., 2019], as the additional-validation data to test eq. (12) on an additional data.

According to the biological mechanism of neoepitopes to generate an immune response: for a given pair of peptide and MHC, the formation and stabilization of the corresponding peptide-MHC (pMHC) respectively requires strong and stable binding between the peptide and the MHC (i.e., high pMHC binding affinity and pMHC binding stability), the presence of pMHC on its cell surface requires abundant expression of the gene generating the peptide (i.e., high tumor abundance), and the recognition of the pMHC by the host immune system requires the pMHC to be either different from other pMHCs (i.e., of low agretopicity) or similar to the pMHCs of some known pathogens (i.e., of high foreignness) [Kindt et al., 2007, chapters 7-10]. Indeed, it has been found that pMHC binding affinity, pMHC binding stability, tumor abundance, agretopicity and foreignness constitute five key features to predict whether a given neoepitope candidate is immunogenic [Wells et al., 2020]. Hence, we used these five features to propose the general form of eq. (12) to predict the probability that a neoepitope candidate is immunogenic (i.e., positive as validated by multimer-staining assay).

Let DtnQUAL(*x*) be the allele quality of the mutation that is transcribed and translated into the peptide of *x*, where the allele quality is defined by the QUAL column in the VCF file format specification [Danecek et al., 2011] and computed by UVC in tumor-normal-paired mode from DNA-seq data [Zhao et al., 2022]. Let DtAD(*x*), DtAF(*x*) and DnAF(*x*) be the tumor allele depth, tumor allele fraction and normal allele fraction (of the mutation transcribed and translated into the peptide of *x*) computed by UVC from DNA-seq data, respectively [Zhao et al., 2022]. Let RtAD(*x*) and RtAF(*x*) be the tumor mutant allele depth and tumor mutant allele fraction (of the mutation translated into the peptide of *x*) computed by UVC from RNA-seq data, respectively [Zhao et al., 2022]. Let Affinity(*x*) and BindLevel(*x*) be the binding affinity (with the unit of nanomolar of dissociation constant) and binding level of the pMHC *x* computed by netMHCpan, respectively [Reynisson et al., 2020]. Let Stability(*x*) be the binding stability (with the unit of hour) of the pMHC *x* computed by netMHCstabpan [Rasmussen et al., 2016]. Let Agretopicity(*x*) be defined and computed as Affinity(*x*)*/* Affinity(*w*) where *w* is the wild-type peptide corresponding to the mutant peptide of *x* and where *x* is also the neoepitope candidate of interest [Wells et al., 2020]. Let Foreignness(*x*) be defined as the probability of the recognition of the pMHC *x* by a random T-cell receptor (TCR) and computed in exactly the same way as the antigen.garnish R package [Richman et al., 2019, Wells et al., 2020]. Let Abundance(*x*) be the tumor abundance computed by using Kallisto [Bray et al., 2016] to obtain the transcript per million (TPM) of each gene transcript and then by summing up the TPMs of all transcripts that carry the mutation generating the peptide of *x*. Then, for each pMHC (i.e., neoepitope candidate) *x ∈ X*, NeoHeadHunter computes the probability *P* that the peptide of *x* is an immunogenic neoepitope as follows:

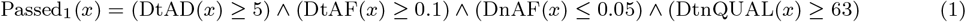

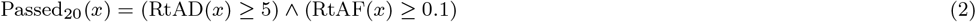

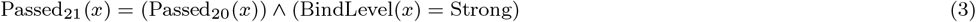

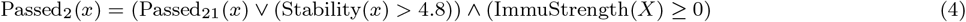

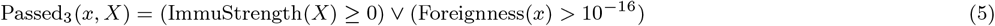

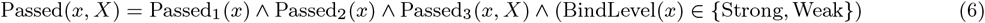

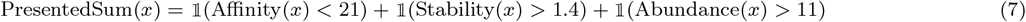

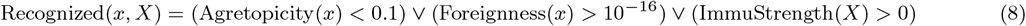

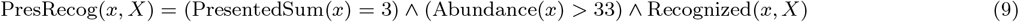

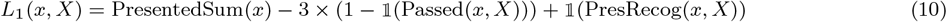

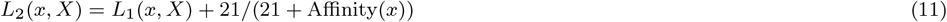

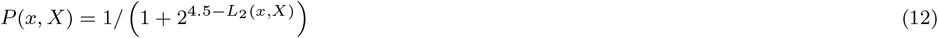

where

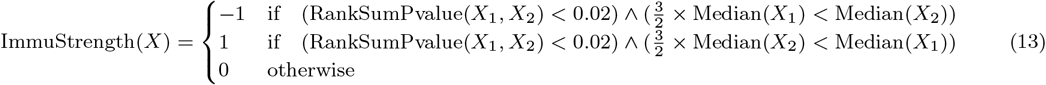

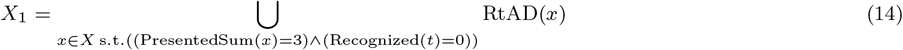

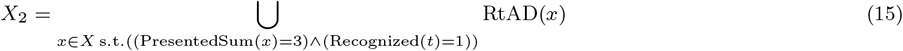

and where PresentedSum(*x*) *∈* {0, 1, 2, 3} is the strength that the peptide is presented by the MHC to the surface of cancer cells for the pMHC *x*, Recognized(*x*) *∈* {0, 1} indicates whether the peptide of *x* is foreign to the host immune system, PresRecog(*x*) *∈* {0, 1} is the strength that the peptide of *x* is recognized by the T cells of the host immune system as a foreign peptide, ImmuStrength(*X*) *∈ {−*1, 0, 1} is the extent to which the host immune system is capable of recognizing the peptides of *X* generated by the cancer cells of the patient of interest, 𝟙 is the indicator function taking TRUE and FALSE Boolean values as input and respectively returning 1 and 0 as output, RankSumPvalue(*A, B*) computes the pvalue of the Wilcoxon (or equivalently Mann–Whitney) rank-sum test for the two sets of input values *A* and *B*, Median(*A*) computes the median of a set *A* of input values, *∧* is the logical AND operator, *∨* is the logical OR operator, and s. t. denotes “such that” in the set-builder notation.

The thresholds of 5 and 0.1 for both DtAD and RtAD and both DtAF and RtAF in eqs. (1) and (3) were also used by NeoHunter [Ma, 2023] and are reasonable thresholds used to filter out false positive somatic variants [Koboldt et al., 2013]. The threshold of 0.05 for DnAF in eq. (1) was also used by NeoHunter [Ma, 2023] and are reasonable thresholds used to consider the effect of contamination [Cibulskis et al., 2011, Taylor-Weiner et al., 2018]. The threshold of 63 for DtnQUAL in eq. (1) is equivalent to the tumor log-odds score (TLOD) threshold of 6.3 used by Mutect [Cibulskis et al., 2013]. The threshold of 4.8 to rescue immunogenic neoepitopes with Stability in eq. (4) is sufficiently high to filter out a lot of neoepitope candidates: for the five patients with publicly available sequencing data in the TESLA study in the discovery cohort, Stability *>* 4.8 for only 108 out of the 2248 NeoHeadHunterdetected neoepitope candidates that failed the Passed_21_ filter in eq. (3).

The TESLA study established the binding affinity, binding stability, tumor abundance, agretopicity and foreignness thresholds of 34, 1.4, 33, 0.1 and 10^*−*16^ by performing repeated random sampling on the discovery cohort consisting of six patients [Wells et al., 2020]. Thus, we also used these thresholds in eq. (12) with the following two exceptions.

1. The TESLA study and we used netMHCpan versions 4.0 and 4.1b to compute pMHC binding affinity, respectively [Wells et al., 2020], and netMHCpan version 4.1b predicts lower dissociation constants for immunogenic pMHCs compared with netMHCpan version 4.0. Thus, we used a threshold of 21 for pMHC binding affinity.
2. The TESLA study mentioned that a lot of immunogenic pMHCs did not pass the filter with a tumor abundance threshold of 33 transcript-per-million (TPM) in the discovery cohort [Wells et al., 2020]. Thus, we used a threshold of 11 for tumor abundance.

A p-value that is smaller than 0.02 is significant enough to reject the null hypothesis that PresentedSum and Recognized are independent of each other. Moreover, it is unlikely that one group of genes have more than 50% more expression than another group of genes by chance. Therefore, ImmuStrength = 0 for most patients. Indeed, ImmuStrength = 0 for all six patients in the discovery cohort except that ImmuStrength = *−*1 for the patient with ID=2. Hence, we would expect that the immune system of the patient with ID=2 tends to fail to recognize tumor cells, so neoepitopes from this patient supposedly have to be more foreign to be recognized by the T cells of this patient. Indeed, Foreignness = 1 for all the four immunogenic neoepitopes from the patient with ID=2, whereas Foreignness = 1 for only 14 out of the 37 validated neoepitopes in total for patients in the discovery cohort. Therefore, penalizing pMHCs with Foreignness = 0 for a patient with ImmuStrength = *−*1 is justified.

If ImmuStrength(*X*) = 0 and we sort the probabilities generated by eq. (12) in descending order, then we naturally produce a ranking of neoepitope candidates as follow. First, the candidates are grouped into five classes and ranked according to their classes in the following order.

1. The class of presented and recognized candidates with PresentedSum = 3 and Recognized = 1.
2. The class of presented and unrecognized candidates with PresentedSum = 3 and Recognized = 0.
3. The class of candidates satisfying two presentation criteria with PresentedSum = 2.
4. The class of candidates satisfying one presentation criterion with PresentedSum = 1.
5. The class of candidates not satisfying any presentation criteria with PresentedSum = 0.

Then, the candidates within each class are ranked by pMHC binding affinity.

Finally, eq. (12) results from the application of Platt scaling [Platt et al., 1999] to eq. (11) with a scale parameter of ln(2) and with a location parameter of 4.5. The location and scale parameters of ln(2) and 4.5 can well calibrate the probabilities for the discovery cohort (fig. 5), and these two parameters do not affect the ranking of neoepitope candidates. Wells et al. showed that binding affinity, binding stability and tumor abundance are three independent factors that should all be satisfied for the MHC to present the mutant peptide (i.e., neoepitope candidate) to the cancer cell surface [Wells et al., 2020]. Therefore, the ranking produced by the probabilities generated by eq. (12) is reasonable.

## 3. Implementation

Our NeoHeadHunter pipeline uses BWA MEM and STAR to align DNA-seq and RNA-seq reads to the GRCh37 human reference genome, respectively. Then, our pipeline uses UVC to call small variants (i.e., single-nucleotide variants (SNVs) and insertion-deletions (InDels)) from both DNA-seq data and RNA-seq data. The normal DNA-seq data are used as the matched normal for the tumor RNA-seq data because normal RNA-seq data were not generated. Next, our pipeline uses a custom Python script to translate, for each small variant, the REF and ALT genotypes characterized by the absence and presence of the small variant into wild-type and mutant peptides, respectively. While running the above steps, our pipeline concurrently uses Kallisto to estimate the abundance of each RNA transcript and uses Optitype on the tumor RNA-seq data to perform human-leukocyte antigen (HLA) typing which determines the types of major-histocompatibility complex (MHC) molecules generated by the tumor.

After peptide generation, RNA-transcript abundance estimation and HLA typing, our pipeline uses netMHCpan and netMHCstabpan to predict, for each combination of peptide and MHC, the affinity and stability of the binding between the peptide and MHC to form a peptide-MHC (pMHC), respectively.

We implemented the NeoHeadHunter algorithm using the Snakemake [Köster and Rahmann, 2012] workflow language (fig. 2). We used BWA MEM version 0.7.17 [Li, 2013] for DNA-seq alignments, STAR version 2.7.8 [Dobin et al., 2013] for RNA-seq alignments, STAR-Fusion version 1.12.0 [Haas et al., 2017] for RNA fusion detection, ASNEO commit 9f43cff [Zhang et al., 2020] for RNA splicing-variant detection, Kallisto version 0.48.0 [Bray et al., 2016] for RNA-transcript abundance estimation, UVC version 0.14.2.15f4adc and uvc-delins version 0.1.7.30af093 [Zhao et al., 2022] for small-variant calling, Ensembl-VEP version 109.3 [McLaren et al., 2016] for small-variant effect prediction, Optitype version 1.3.2 [Szolek et al., 2014] with sequencing data pre-filtered by bwa mem for HLA typing, NetMHCpan version 4.1b [Reynisson et al., 2020] for peptide-MHC binding affinity prediction and NetMHCstabpan version 1.0 along with NetMHCpan version 2.8 [Rasmussen et al., 2016] for peptide-MHC binding stability prediction. Additionally, we used the CTAT resource library `GRCh37_gencode_v19_CTAT_lib_Mar012021` [Haas et al., 2019] for DNA-seq alignments, RNA-seq alignments and fusion detection.

**Figure 2:**
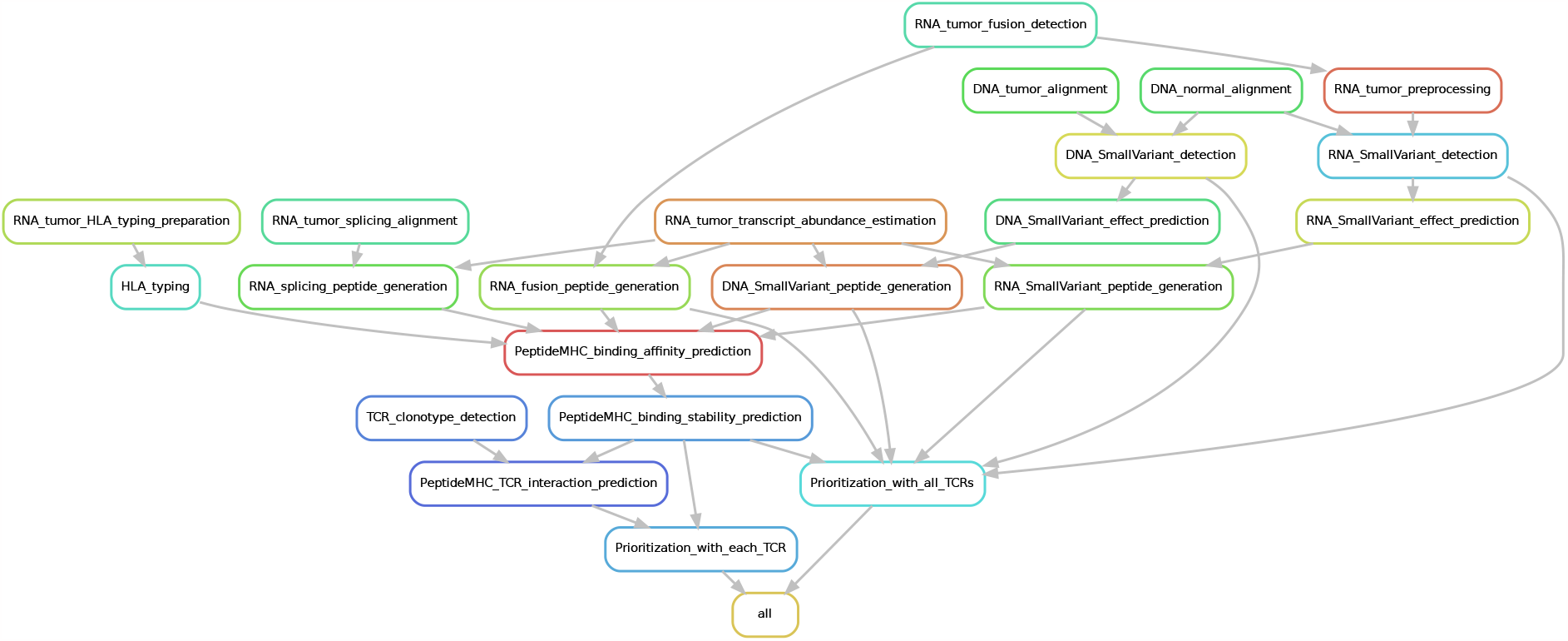
Schematics of the NeoHeadHunter workflow generated by Snakemake [Köster and Rahmann, 2012] and Graphviz [Ellson et al., 2004].

NeoHeadHunter can also run STAR-Fusion [Haas et al., 2017] and ASNEO [Zhang et al., 2020] on the tumor RNA-seq data to generate neoepitope candidates from fusion and splicing variants, respectively. Unfortunately, as far as we know, both validation of immunogenity and report of clinical benefit are absent in the literature for fusion and splicing variants. Therefore, the generation of neoepitope candidates from fusion and splicing variants is incorporated as an optional step in NeoHeadHunter but not described in detail.

NeoHeadHunter can also run MixCR [Bolotin et al., 2015] and ERGO-II [Springer et al., 2021] on the tumor RNA-seq data to prioritize neoepitopes for each specific T-cell receptor (TCR) clonotype. Unfortunately, it has been shown that neoepitope prioritization with respect to each specific TCR clonotype performs much worse than with respect to all TCR clonotypes as an ensemble presumably because TCR clonotypes are too diverse to be captured by a typical sequencing assay [Ma, 2023] and because the interaction between TCR and pMHC is hard to predict (especially for neoepitopes which are absent in the training set, and such neoepitopes constitute the vast majority of all neoepitopes) [Cai et al., 2022, Hudson et al., 2023]. Therefore, TCR-specific prioritization is incorporated as an optional step in NeoHeadHunter but not described in detail.

## 4. Results

The results of detecting and ranking neoepitope candidates on the discovery cohort are shown in section 4.1, except that the TESLA patient with ID=10 is in the discovery cohort but does not generate any result in section 4.1 because the raw sequencing data of this patient are not available. The results of ranking neoepitope candidates on the independent-validation cohort and the additional data are respectively shown in sections 4.2 and 4.3. The results of probabilistically classifying neoepitope candidates on the discovery, validation, and independent-validation cohorts are shown in section 4.4.

### 4.1 Task of detecting and ranking neoepitope candidates for five TESLA patients with raw sequencing data

TESLA published the raw sequencing data for five patients (with the patient IDs of 1, 2, 3, 12 and 16) and used top-twenty immunogenic fraction (TTIF), fraction ranked (FR) and area under the precision-recall curve (AUPRC) as the metrics in their evaluation. Therefore, we also used TTIF, FR and AUPRC to evaluate the performance of NeoHeadHunter and other algorithms on these five patients. Moreover, we used TFA-mean, defined as the average of TTIF, FR and AUPRC, as an additional summary metric in our evaluation.

NeoHeadHunter overall performs the best in terms of TTIF, FR and AUPRC (fig. 3). In terms of TFA-mean, NeoHeadHunter performs the best for each patient except that the owl algorithm performs better than NeoHeadHunter for the patient with ID=16 (fig. 3). We manually reviewed all variant candidates generated by UVC and found that, for the patient with ID=16, a double-nucleotide variant (DNV) that translates into the peptide `YLNEAMFNFV` was presumably called incorrectly as a singlenucleotide variant (SNV) that translates into the peptide `YLNEAVFNFV`, so the immunogenic (i.e., positive as validated by multimer-staining assay) neoepitope `YLNEAVFNFV` is actually generated by a variantcalling error and should be replaced by `YLNEAMFNFV`. Thus, this error can explain the under-performance of NeoHeadHunter for the patient with ID=16.

**Figure 3:**
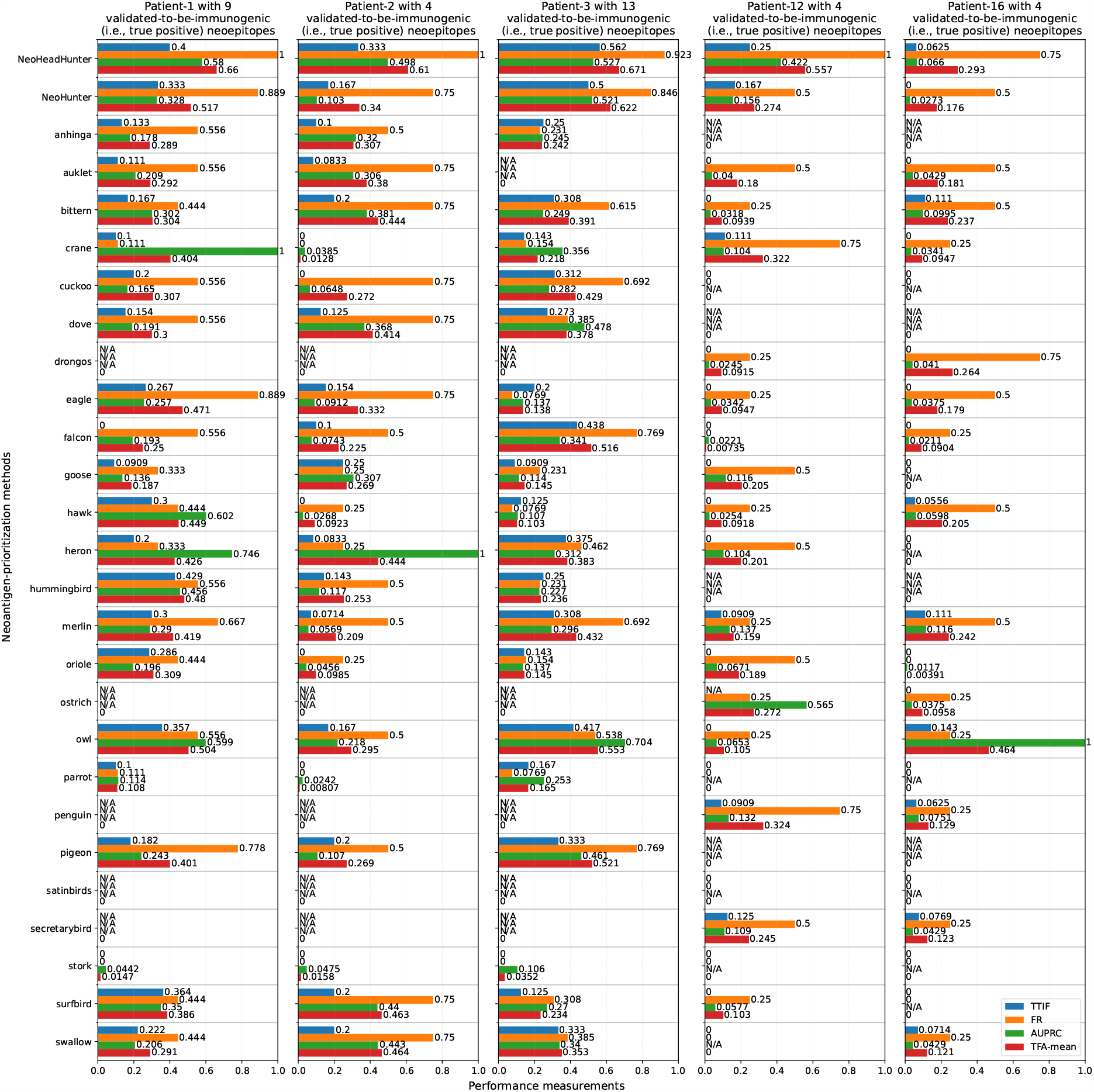
Performance of NeoHeadHunter and other methods on the TESLA dataset for finding and ranking neoepitope candidates [Wells et al., 2020]. TTIF stands for top-twenty immunogenic fraction. FR stands for fraction ranked. AUPRC stands for area under the precision-recall curve. TFA-mean stands for the average of TTIF, FR and AUPRC.

UVC is able to call the SNV (that Mutect2 [Benjamin et al., 2019] missed due to low mapping quality) that translates into the immunogenic peptide `FLNCDIMLGV`. Additionally, UVC is able to call the SNV responsible for generating the immunogenic peptide `KIVEMSTSK` using a combination of RNAseq tumor data and DNA-seq normal data. In sum, UVC is able to call all variants that translate into the immunogenic peptides, with the exception that UVC missed a single-nucleotide variant (SNV) that translates into the peptide `YLNEAVFNFV`, but further manual review shows that this SNV should be called as a double-nucleotide variant (DNV) that translates into the peptide `YLNEAMFNFV`. Thus, after excluding the erroneous peptide YLNEAVFNFV, we observed that NeoHeadHunter detected all the 33 immunogenic peptides in total from the five TESLA patients with the IDs of 1, 2, 3, 12 and 16. Hence, FR and detection rate are slightly improved by the use of UVC to call SNVs and InDels. The use of eq. (11) to rank neoepitope candidates results in high performance in terms of AUPRC and TTIF because candidates that are more likely to be immunogenic are ranked before the other candidates. The use of eq. (12) is able to predict, for each neoepitope candidate, its probability of being immunogenic, resulting in the probabilistic classification of neoepitope candidates.

Moreover, we compared the performance of NeoHeadHunter with TSNAD v2.0 [Zhou et al., 2021] using the evaluation criteria that TSNAD v2.0 used. More specifically, the number of pMHC pairs that were tested for immunogenity (#Tested), the number of pMHC pairs that were tested to be immunogenic (#True) and the ratio of #True to #Tested (Accuracy) are listed in table 1 when selecting pMHC pairs ranked top 10, 20, 30, 40 and 50 as high-confidence neoepitopes, respectively. NeoHeadHunter offers significant improvement compared with TSNAD v2.0 on the five TESLA patients with the IDs of 1, 2, 3, 12 and 16 (table 1).

**Table 1:** The performance of TSNAD v2.0 and NeoHeadHunter under different selection criteria [Zhou et al., 2021, Wells et al., 2020]. The column #Tested denotes the number of peptide-MHC (pMHC) pairs that were subject to validation, where MHC stands for major histocompatibility complex. The column #True denotes the number of pMHC pairs that are immunogenic (i.e., positive as validated by multimer-staining assay). The column Accuracy is defined as #True divided by #Tested.

| Selection<br>criterion | TSNAD v2.0 with GRCh38 |  |  | TSNAD v2.0 with GRCh37 |  |  | NeoHeadHunter |  |  |
| --- | --- | --- | --- | --- | --- | --- | --- | --- | --- |
|  | #Tested | #True | Accuracy | #Tested | #True | Accuracy | #Tested | #True | Accuracy |
| Top10 | 18 | 2 | 11.1% | 17 | 2 | 11.8% | 37 | 13 | 35.1% |
| Top20 | 23 | 5 | 21.7% | 30 | 5 | 16.7% | 63 | 21 | 33.3% |
| Top30 | 27 | 5 | 18.5% | 35 | 5 | 14.3% | 87 | 25 | 28.7% |
| Top40 | 33 | 6 | 18.2% | 43 | 5 | 11.6% | 103 | 27 | 26.2% |
| Top50 | 40 | 6 | 15.0% | 49 | 6 | 12.2% | 122 | 29 | 23.8% |

### 4.2 Task of ranking neoepitope candidates on three NSCLC patients with MuPeXI-annotated neoepitope candidates

Bjerregaard et al. ran MuPeXI to generated a list of 190 neoepitope candidates in total for three NSCLC patients, and exactly eight of these 190 candidates are immunogenic [Bjerregaard et al., 2017, McGranahan et al., 2016, Bentzen et al., 2016]. For each of these NSCLC patients, MuPeXI also provided a ranking of these candidates such that a higher rank implies a lower priority. We used NeoHeadHunter to re-rank these candidates for each patient. For each of these eight immunogenic candidates, NeoHeadHunter always provides a rank that is not greater than the rank provided by MuPeXI (table 2). Hence, this independent dataset shows that NeoHeadHunter performs well for ranking neoepitope candidates because NeoHeadHunter consistently ranks immunogenic candidates before the other candidates.

**Table 2:**
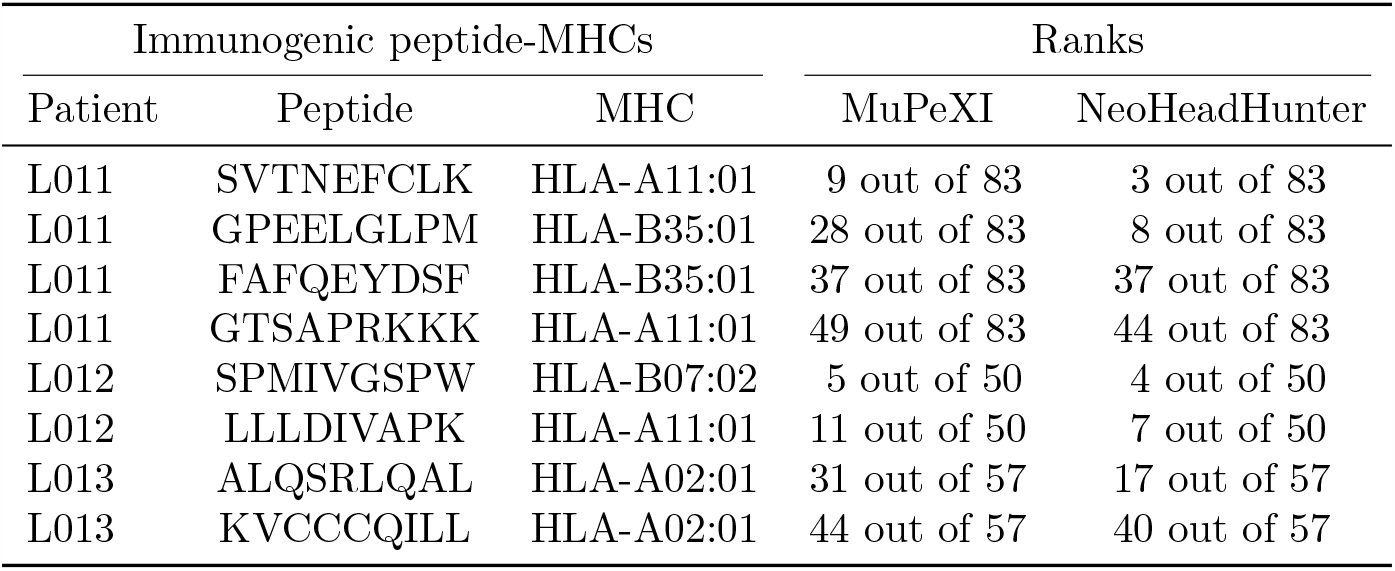
Performance of MuPeXI and NeoHeadHunter for ranking immunogenic (i.e., positive as validated by multimer-staining assay) neoepitope candidates before the other candidates. The candidates are detected by MuPeXI from three non-small-cell lung cancer (NSCLC) patients [Bjerregaard et al., 2017]. Lower ranks imply higher priorities. MHC stands for major histocompatibility complex.

| Immunogenic peptide-MHCs |  |  | Ranks |  |
| --- | --- | --- | --- | --- |
| Patient | Peptide | MHC | MuPeXI | NeoHeadHunter |
| L011 | SVTNEFCLK | HLA-A11:01 | 9 out of 83 | 3 out of 83 |
| L011 | GPEELGLPM | HLA-B35:01 | 28 out of 83 | 8 out of 83 |
| L011 | FAFQEYDSF | HLA-B35:01 | 37 out of 83 | 37 out of 83 |
| L011 | GTSAPRKKK | HLA-A11:01 | 49 out of 83 | 44 out of 83 |
| L012 | SPMIVGSPW | HLA-B07:02 | 5 out of 50 | 4 out of 50 |
| L012 | LLLDIVAPK | HLA-A11:01 | 11 out of 50 | 7 out of 50 |
| L013 | ALQSRLQAL | HLA-A02:01 | 31 out of 57 | 17 out of 57 |
| L013 | KVCCCQILL | HLA-A02:01 | 44 out of 57 | 40 out of 57 |

### 4.3 Task of ranking neoepitope candidates on a manually curated set of neoepitope candidates

Wu et al. showed that DeepHLApan performs well compared with other competing algorithms on the dataset curated by Koşaloğlu-Yalçın et al. [2018] [Wu et al., 2019]. Hence, we evaluated the performance of DeepHLApan and NeoHeadHunter on this dataset. Our evaluation shows that NeoHeadHunter performs better than DeepHLApan in terms of area under the curve of receiver operating characteristic (AUC-ROC). (fig. 4). Hence, this independent dataset shows that NeoHeadHunter performs well for ranking neoepitope candidates.

**Figure 4:**
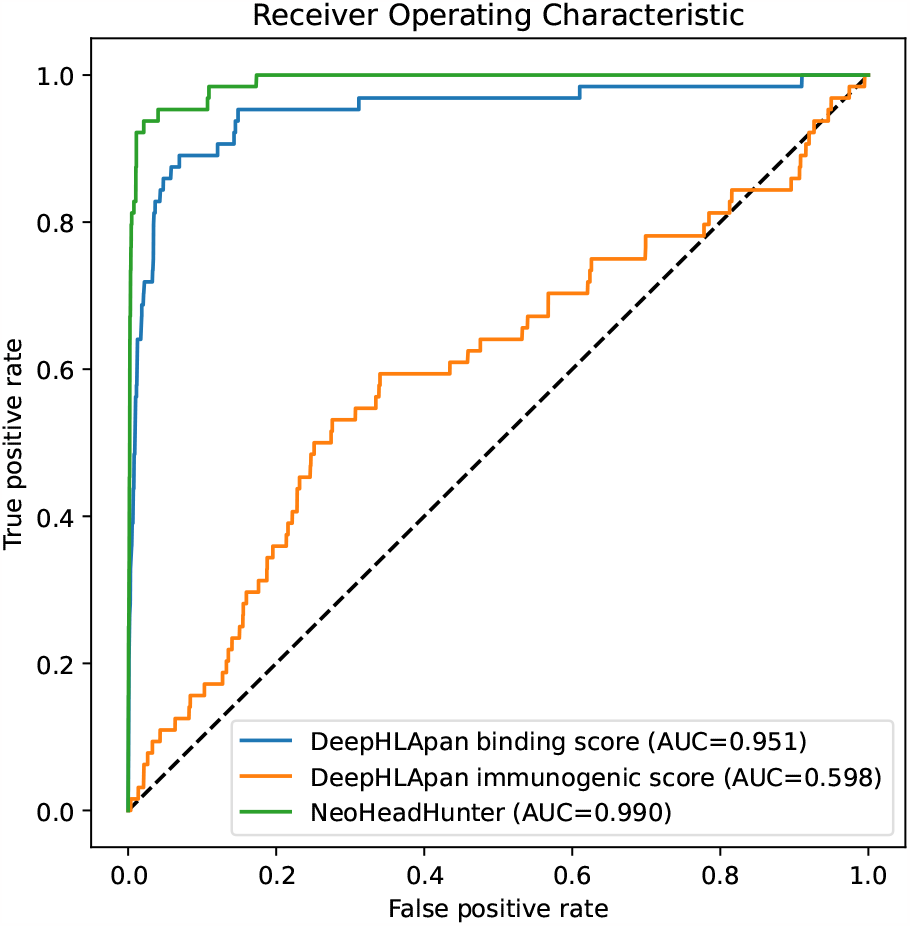
Area under the curve of receiver operating characteristic (AUC-ROC) for DeepHLApan [Wu et al., 2019] and NeoHeadHunter evaluated on the dataset curated by Koşaloğlu-Yalçın et al. [2018]. The curated dataset consists of 64 immunogenic (i.e., positive as validated by an immuno-assay) neoepitopes extracted from the literature and 6400 random peptide-MHC pairs generated based on mutation data extracted from The Cancer Genome Atlas (TCGA) database. MHC, major-histocompatibility complex.

**Figure 5:**
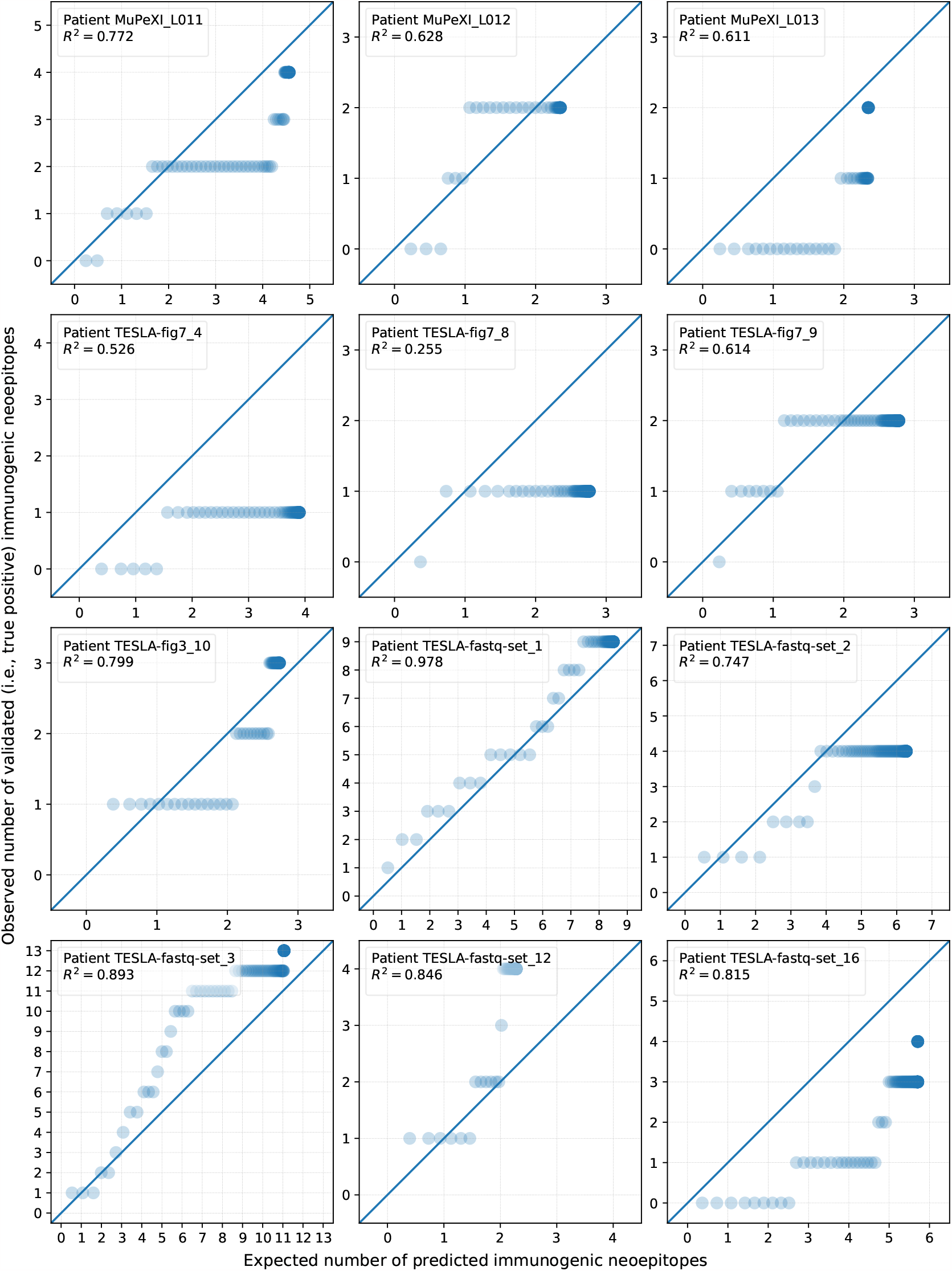
Performance of NeoHeadHunter on the TESLA [Wells et al., 2020] and MuPeXI-annotated NSCLC [Bjerregaard et al., 2017, McGranahan et al., 2016, Bentzen et al., 2016] datasets for probabilistically classifying neoepitope candidates. For each patient, the candidates are sorted in descending order by their predicted probabilities of being immunogenic (i.e., positive as validated by multimer-staining assay), and cumulative summations of predicted probability and actual binary outcomes are performed on the sorted candidates. The cumulative sums of predicted probabilities and actual binary outcomes are shown on the x-axis and y-axis, respectively.

### 4.4 Task of predicting immunogenity probabilities of neoepitope candidates for five TESLA patients with raw sequencing data, four TESLA patients with TESLA-annotated neoepitope candidates, and three NSCLC patients with MuPeXI-annotated neoepitope candidates

TESLA also published the lists of neoepitope candidates for four patients with the IDs of 4, 8, 9 and 10. Therefore, we used observed versus expected cumulative number of immunogenic neoepitopes to evaluate the probabilistic classification of NeoHeadHunter on the 12 patients with either raw sequencing data or lists of neoepitope candidates. Our cumulative number is computed in the same way as the one defined by Arrieta-Ibarra et al. [Arrieta-Ibarra et al., 2022] except that

1. we used cumulative sum instead of cumulative average because the total effectiveness of the neoepitope-based vaccine is of more interest and that

2. we sorted the probabilities in descending order instead of ascending order because neoepitope candidates with higher probabilities of immunogenicity are of more interest.

Because NeoHeadHunter is the only algorithm that can probabilistically classify neoepitope candidates, we did not compare NeoHeadHunter with any other algorithms for the task of probabilistic classification. The theoretical probabilities of immunogenicity predicted by NeoHeadHunter closely match the empirical probabilities observed from validation (fig. 5).

Only very few neoepitope features are available for the four TESLA patients with the IDs of 4, 8, 9 and 10. Consequently, we are not able to evaluate the ranking of NeoHeadHunter compared with other algorithms. Nevertheless, for these four patients, NeoHeadHunter also ranked the very few immunogenic neoepitope candidates before the non-immunogenic ones (fig. 5). Thus, NeoHeadHunter is supposed to performs well for ranking neoepitope candidates for these four patients.

### 4.5 Evaluation detail

Top-twenty immunogenic fraction (TTIF) is the ratio of the number of top-20 ranked pMHCs with detected immunogenicity to the total number of top-20 pMHCs which were tested for immunogenicity [Wells et al., 2020]. Fraction ranked (FR) is the fraction of all peptides with detected immunogenicity for a particular subject that were included in the top 100 ranked pMHC by a participant [Wells et al., 2020]. Area under the precision-recall curve (AUPRC) was calculated using the pr.curve function in the PRROC package [Grau et al., 2015] with default parameters [Wells et al., 2020]. TTIF, FR and AUPRC were all calculated using the R code at https://github.com/ParkerICI/tesla/blob/master/performance-metric-functions.R [Wells et al., 2020]. TFA-mean is the average of TTIF, FR and AUPRC.

We assessed the performance of MuPeXI [Bjerregaard et al., 2017] using the strategy of ranking neoepitope candidates by the `priority_Score` generated by MuPeXI for all applicable patients. The TTIF and FR of MuPeXI is zero for all the five patients (i.e., patients with the IDs of 1, 2, 3, 12 and 16) with publicly available sequencing data. Hence, the AUPRC of MuPeXI is also very close to zero for all these five patients. Thus, the TTIF, FR, AUPRC and TFA-mean of MuPeXI are not shown for these five patients (fig. 3).

For the four TESLA patients with TESLA-annotated neoepitope candidates (with the IDs of 4, 8, 9 and 10) and the three NSCLC patients with MuPeXI-annotated neoepitope candidates (with the IDs L011, L012 and L013), we let Passed_1_ and Passed_20_ return the TRUE Boolean value because the values of the features required to compute Passed_1_ and Passed_21_ are missing for these seven patients, and we ran blast version 2.12.0+ [Altschul et al., 1990] with the parameters “-word size 3 -evalue 1e8” against the peptide database of VEP version 109 [McLaren et al., 2016] to find the wild-type peptide of each mutant peptide in order to calculate agretopicity. For each neoepitope candidate, we let Abundance = 10 if Abundance is not available because the median Abundance is approximately 10 in the TESLA dataset [Wells et al., 2020, figs. 3 and 7].

In our evaluation, we ran NeoHeadHunter commit 078d14e, NeoHunter commit 79e38d9 [Ma, 2023], MuPeXI commit 1970565 [Bjerregaard et al., 2017] and DeepHLApan version 18.09 [Wu et al., 2019]. In fig. 3, table 1 and table 2, we directly showed the previously published performances of the TESLA-evaluated algorithms (i.e., anhinga, auklet, …, swallow) [Wells et al., 2020, table S5], TSNAD 2.0 [Zhou et al., 2021, table 4] and MuPeXI 1.1 [Bjerregaard et al., 2017, table S2], respectively.

Tools such as pVAC-seq [Hundal et al., 2016], NeoANT-HILL [Coelho et al., 2020] and NeoFox [Lang et al., 2021] are known to be unable to produce any ranking of neoepitope candidates, presumably because their design philosophy is to annotate each neoepitope candidate with a lot of features and to expect the user to use these features along with his/her domain knowledge about neoantigens to manually rank these neoepitope candidates (e.g., by eliminating all candidates with netMHCpan-predicted binding affinity of more than 500nM). Therefore, such tools cannot be evaluated with the ranking-based performance metrics including TTIF, FR, AUPRC, TFA-mean (fig. 3), Top10, Top20, Top30, Top40, Top50 (table 1) and ranks (table 2). Hence, we did not evaluate such tools because they were not designed to produce any ranking of neoepitope candidates.

## 5 Discussion

We designed and developed NeoHeadHunter, an algorithm and pipeline to detect, rank, and probabilistically classify neoepitope candidates. NeoHeadHunter can detect all immunogenic neoepitopes, can accurately rank neoepitope candidates, and can accurately estimate the probability that each neoepitope candidate is immunogenic (i.e., positive as validated by multimer-staining assay). For each patient, the expected number of immunogenic neoepitopes (tumor neoepitope burden, TNB), defined as the sum of all such NeoHeadHunter-estimated probabilities of all neoepitope candidates, may be positively correlated with the overall effectiveness of the personalized cancer vaccine given to this patient. It is likely that TNB can predict the response to therapy based on personalized cancer vaccine, similar to the way that tumor mutation burden (TMB) can predict the response to therapy based on immune checkpoint inhibitors [Jardim et al., 2021].

Equation (9) implies that Recognized is always TRUE if ImmuStrength = 1, which conforms to our intuition that Agretopicity and Foreignness become less important if ImmuStrength becomes high. Unfortunately, we did not find any patient with ImmuStrength = 1. Therefore, the effect of ImmuStrength on Recognized needs to be validated by patients with ImmuStrength = 1 even though finding such patients may not be easy. Additionally, in real clinical settings, we can estimate ImmuStrength in eq. (13) from other patient-related features such as age and gender so that the scarcity of presented peptides does not hinder the estimation of ImmuStrength.

For all patients that we used in our evaluation, a neoepitope candidate is labeled to be immunogenic if tested positive by the corresponding multimer-staining assay, and vice versa. Thus, eq. (12) with a different location parameter can possibly fit better to the ground-truth of immunogenicity established by another immuno-assay.

## 6 Conclusions

NeoHeadHunter is an easy-to-use algorithm and pipeline to prioritize neoepitopes. NeoHeadHunter has the ability to sensitively detect, accurately rank, and probabilistically classify neoepitope candidates. Thus, NeoHeadHunter is able to increase the effectiveness of personalized cancer vaccine.

## Supporting information

Results generated by NeoHeadHunter with validation

## 7 Availability and requirements

**Project name:** NeoHeadHunter

**Project home page:** https://github.com/XuegongLab/neoheadhunter

**Operating system(s):** Linux and WSL (Windows Subsystem for Linux)

**Programming language:** Snakemake

**Other requirements:** Python 3 with Bioconda

**License:** APACHE-II

**Any restrictions to use by non-academics:** the third-party software packages UVC, netMHCpan, netMHCstabpan and MixCR require licensing for non-academic use.

## 8 Declarations

### 8.1 Ethics approval and consent to participate

Not applicable

### 8.2 Consent for publication

Not applicable

### 8.3 Availability of data and materials

All data used in this manuscript are available at https://www.synapse.org/#!Synapse:syn21048999, Wells et al. [2020, Tables S4, S5 and S7] and Bjerregaard et al. [2017, Table S2].

### 8.4 Competing interests

The authors declare no competing interests.

## 8.5 Funding

Xiaofei Zhao is supported by the Chinese Government Scholarship.

## 8.6 Authors’ contributions

Not applicable

## 8.7 Acknowledgements

We would like to thank Dr Xuegong Zhang and Dr Lei Wei for providing some comments on how to revise this manuscript.

## Notes

### Competing Interest Statement

The authors have declared no competing interest.

